# Quantification of Wnt3a, Wnt5a and Wnt16 Binding to Multiple Frizzleds Under Physiological Conditions using NanoBit/BRET

**DOI:** 10.1101/2025.04.10.648195

**Authors:** Janine Wesslowski, Sadia Safi, Michelle Rottmann, Melanie Rothley, Gary Davidson

**Affiliations:** Institute of Biological and Chemical Systems – Functional Molecular Systems (IBCS-FMS), Karlsruhe Institute of Technology (KIT), Karlsruhe, Germany

**Keywords:** Wnt Signaling, Ligand-Receptor binding, Wnt-FZD binding, NanoBiT/BRET, HiBiT, Frizzled, GFP-Wnt, HiBiT-FZD, LRP6, LRP6-Wnt-FZD complex

## Abstract

Upon engagement of one of the 19 secreted Wnt signalling proteins with one of the 10 Frizzled transmembrane Wnt receptors (FZD_1-10_), a wide variety of cellular Wnt signalling responses can be elicited, the selectivity of which depends on: 1) the specific Wnt-FZD pairing, 2) the participation of Wnt co-receptors, and 3) the cellular context. Co-receptors play a pivotal role in guiding the specificity of Wnt signaling, most notably between β-catenin dependent and independent pathways, where co-receptors such as LRP5/6 and ROR1/2 / PTK7 play major roles, respectively. It remains less understood how specific Wnt/FZD combinations contribute to the selectivity of downstream Wnt signaling and we lack accurate comparative data on their binding properties under physiological conditions. Here, using fluorescently-tagged Wnt3a, Wnt5a and Wnt16 proteins and cell lines expressing HiBiT-tagged Frizzled, we build on our ongoing efforts to provide a complete overview of the biophysical properties of all Wnt/FZD interactions using full-length proteins. Our real-time NanoBRET analysis using living cells expressing low receptor levels provides more accurate quantification of binding and will help us understand how these binary engagements control Wnt signaling outputs. We also provide evidence that LRP6 regulates the binding affinity of Wnt/FZD interactions in the trimeric Wnt-FZD-LRP6 complex.

## Introduction

Wnt signaling is initiated and controlled at the cell membrane through the interaction of 19 secreted Wnt lipo-glycoproteins (ligands), 10 principal Wnt receptors called Frizzled (FZD_1-10_) and several Wnt co-receptors such as LRP5/6, ROR1/2, PTK7 and RYK [1-6]. The intercellular Wnt signals transduced from this large repertoire of possible ligand-receptor interactions are diverse and regulate a multitude of developmental processes as well as tissue homeostasis in adults [1, 2, 7]. The biophysical mechanisms that select pairing of ligand-receptor interactions must therefore play a key role in specifying Wnt signalling and much effort has been placed on studying the relative strengths of Wnt-FZD interactions. The first evidence that differential Wnt/FZD binding affinities regulate Wnt signaling emerged from *Drosophila* studies, where the stronger signaling of Dfz2 compared to Dfz1 correlated with a tenfold increase in binding affinity for Wg [8]. Additionally, an earlier study revealed that two mammalian FZD family members (FZD3 and 6) could not bind Wg, highlighting the divergent nature of Wnt/FZD interactions and the non-canonical properties associated with these particular two Frizzled’s [9]. Indeed, some Wnts preferentially activate non-canonical (β-catenin independent) or canonical (β-catenin dependent) signaling. Canonical ligands include Wnt1, Wnt3a, Wnt7a and Wnt8, whereas non-canonical ligands include Wnt2, Wnt4, Wnt5a, Wnt7b and Wnt11 [1, 10, 11]. It is important to note, however, that there are no strict preferences for individual Wnts because promiscuous effects arise due to different cellular contexts, e.g., because of differences in availability of Wnt receptors and co-receptors [12]. Variations of Wnt signalling beyond β-catenin dependent/independent pathways also exist, further diversifying the biological functions of this essential cellular communication network [1, 13-16].

The ten Frizzled proteins are regarded as the principle Wnt receptors and are instrumental for transduction of all known Wnt signalling events whereas the Wnt co-receptors LRP5/6 appear to be essential only for β-catenin dependent as well as Wnt/STOP signalling [6, 13, 17-19]. With such a large repertoire of ligand-receptor interactions controlling Wnt signaling, a systematic approach to accurately quantify them under native conditions is important and we have made considerable progress in this endeavour over the past few years [20-23]. Here, the term native conditions corresponds to the study of full-length proteins in live cells, using real-time measurements. Indeed, we have demonstrated that eGFP-Wnt3a conditioned medium (CM), when applied to HEK293TA cells overexpressing HiBiT-FZD_1-10_ constructs, allows comparative quantification of binding affinities and kinetics [21]. One downside of this study, however, was the use of HEK293A cells overexpressing high levels of the HiBiT-FZDs, which underrepresents the true binding affinities [20, 23]. Here, we report on our ongoing efforts to measure a wider range of Wnt-FZD interactions by NanoBiT/BRET-based assays, using physiologically relevant receptor expression levels of full-length proteins. We have built on our ability to prepare full-length fluorescently tagged Wnt proteins as well as full-length HiBiT tagged Frizzleds for real-time analysis in living cells. We show that a wide range of HiBiT-tagged FZD proteins, when expressed at low levels in living cells provides a reliable comparison of binding characteristics for up to 3 different Wnt signaling proteins, Wnt3a, Wnt5a and Wnt16. Additionally, we demonstrate that the affinity of Wnt/FZD interactions are regulated by LRP6 in a manner that correlates with the latter’s signaling ability, which may provide some mechanistic insight of how Wnt signalosomes are formed.

## Results

### Fluorescent Tagging of Wnt5a and Wnt16 preserves secretion and signaling

Wnts are secreted lipoproteins that are difficult to purify in an active state and their tagging is challenging, especially larger fluorescent moieties. Wnt3a and Wnt5a were the first to be purified from conditioned medium [12, 24] and more recently, Wnt3a has been successfully fluorescently tagged with only partial loss of signaling activity [22, 25].

To expand on our analysis of Wnt/FZD interactions using NanoBiT/BRET [20, 21], we added an N-terminal eGFP to eleven Wnt proteins (Wnt1, Wnt2b, Wnt5, Wnt6, Wnt7a, Wnt8a, Wnt9a, Wnt10a, Wnt10b, Wnt11 and Wnt16), following the same principle as we described previously for Wnt3a [22]. Barring eGFP-Wnt11, all fusion proteins were secreted and detected in the medium (Supplementary Figure 1A, see M&M for details). Co-expression of the Wnt binding protein Afamin [26] further increased the amounts detected in the medium, especially for eGFP-Wnt6, -7a, -9a, -10a and -10b. Furthermore, Afamin co-expression enabled secretion of Wnt11 into the medium (Supplementary Figure 1A). We therefore routinely included Afamin co-expression for the preparation of condition medium (CM) containing eGFP-Wnt.

Using confocal microscopy, we next tested the ability of the fluorescently tagged Wnt proteins to associate with NCI-H1703 cells stably expressing HiBiT-FZD_5_ (Figure 1A). Upon addition of CM, all, apart from Wnt11, showed specific membrane association to varying degrees, with Wnt3a and Wnt5a presenting the strongest binding. Interestingly, only Wnt3a, Wnt5a and Wnt16 presented a relatively uniform membrane association, with Wnt1, -2b, -6, -7a, -8a, -9a, -10a and -10b displaying only partial binding to cell membranes. Indeed, eGFP-Wnt3a, -5a and -16 displayed the most consistent association to a variety of FZD expressing cells (Supplementary Figure 1B). Reporter assays for β-catenin-dependent (TOPFLASH) and β-catenin-independent (ATF2, NFAT and AP1) Wnt signaling confirmed that eGFP-Wnt3a, eGFP-5a and eGFP-16 retain functional activity (Figure 1B) and CM for these were prepared using Expi293 suspension cells (see M&M for details). Western Blot analysis of the highest protein concentrations used in the NanoBiT/BRET assay confirmed that there was no obvious degradation of the fusion proteins and we obtained precise concentrations for each: eGFP-Wnt3a 15,2 nM; eGFP-Wnt5a 17,3 nM; eGFP-Wnt16 24 nM (Figure 1C).

**Figure 1.**
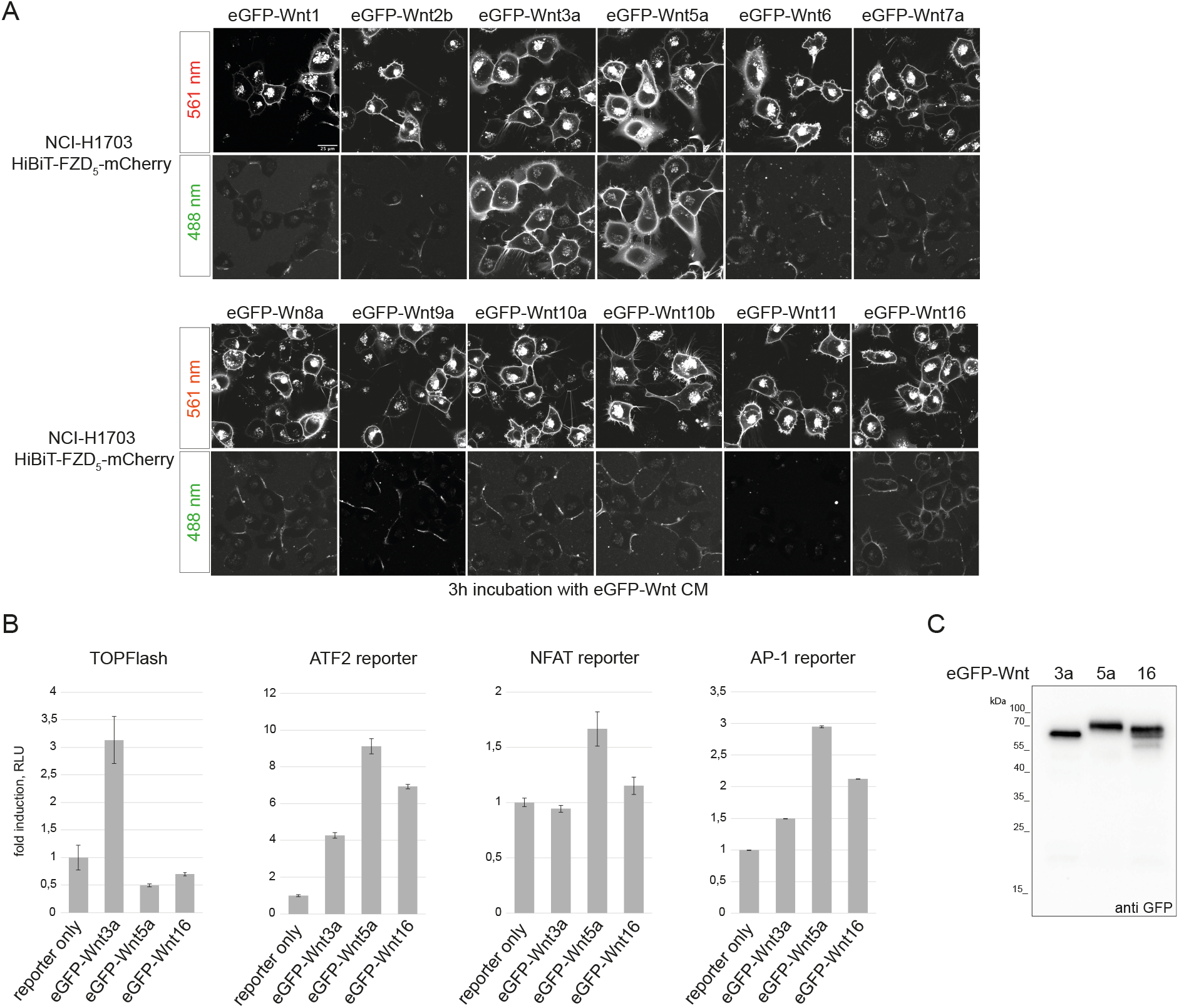
Generation of eGFP-Wnt3a, -5a, -16 CM and U-2 OS cells stably expressing HiBiT-FZD_1-10_-mCherry. **A)** Laser scanning confocal microscopy images of NCI-H1703 cells with stable integration of mouse HiBiT-FZD_5_, incubated for 3 h with the indicated eGFP-Wnt CM derived from Expi293™ suspension cells. **B)** Reporter gene assay using HEK293T cells transfected with TOPFlash reporter to measure the activation of canonical Wnt pathway and ATF2, NFAT, AP1 luciferase reporter to detect β-catenin independent Wnt pathway activation. Cells were treated with eGFP-Wnt3a, eGFP-Wnt5a and eGFP-Wnt16 CM for 24 h. Error bars represent means ± S.D. from 4 independent samples. Experiments were performed 3 times with similar results. **C)** Western Blot using anti GFP antibody, showing presence of soluble eGFP-Wnt3a, eGFP-Wnt5a and eGFP-Wnt16 in conditioned medium (CM) derived from Expi293™ suspension cells, representing the highest concentration used for NanoBiT/BRET assay; 15,2 nM for eGFP-Wnt3a, 17,3 nM for eGFP-Wnt5a and 24 nM for eGFP-Wnt16.

### Generation of HiBiT-FZD stable cell lines

We have previously shown that transient transfection of HiBiT-FZD in HEK293T cells allows reliable measurement of eGFP-Wnt3a binding [21]. To obtain a more physiologically relevant cell culture-based system for measuring Wnt-FZD interactions we used HCT116, NCI-H1703 and U-2 OS cells to generate stable cell lines with low-level expression of HiBiT-FZD_1_. We were unable to generate stable, low HiBiT-FZD_1_ expressing NCI-1703 cell lines, but the larger U-2 OS cells and smaller HCT116 cells were more amenable (Supplementary Figure 2A). Although the average number of receptors per cell for U-2 OS and HCT116 cells are similar, receptor density is significantly lower on U-2 OS due to their larger size (5-to 10-fold larger) and, correspondingly, a weaker fluorescence signal at the cell surface is detected in U-2 OS cells (Supplementary Figure 2B). Next, to study the effect of receptor levels on eGFP-Wnt3a binding using NanoBiT/BRET assays, three different stable U-2 OS cell populations with varying receptor densities were prepared (Figure 2A,B). Stable cell lines expressing lower HiBiT-FZD_1_ levels display a higher binding affinity for eGFP-Wnt3a, which is in line with our previous findings [23] (Figure 2C). This is likely due to a more physiological environment for ligand-receptor binding, minimizing out-titration of endogenous cellular components that play roles in Wnt receptor engagement. Expectedly, a higher standard deviation of measurements is seen in U-2 OS cells expressing lower receptor levels (Figure 2C, compare error bars in binding curves). For all subsequent analysis, we employed FACS-sorted, stable U-2 OS cell lines expressing low/medium levels of HiBiT-FZD_1-10_-mCherry, corresponding to receptor densities of between 500 and 2000 per cell (Figure 2A, Supplementary Figure 2C). The signaling activity of HiBiT-FZD_1-10_-mCherry proteins were tested using TOPFLASH Wnt reporter assays after expressing the corresponding constructs in ΔFZD_1-10GFP-free_ HEK293 cells [22]. No signalling activity was detected for FZD_3_ or FZD_6_ fusions proteins, which is in line with their preference for non-canonical Wnt pathways (Supplementary Figure 2D). FZD_8_ displayed only weak signalling, whereas all others transduced Wnt/β-catenin signaling to a similar degree (Supplementary Figure 2D). Interestingly, although HiBiT-FZD_9_-mCherry showed strong signaling activity in TOPFLASH assays, eGFP-Wnt3a binding was undetectable in HiBiT-FZD_9_-mCherry U-2 OS cells (Supplementary Figure 2C,D).

**Figure 2.**
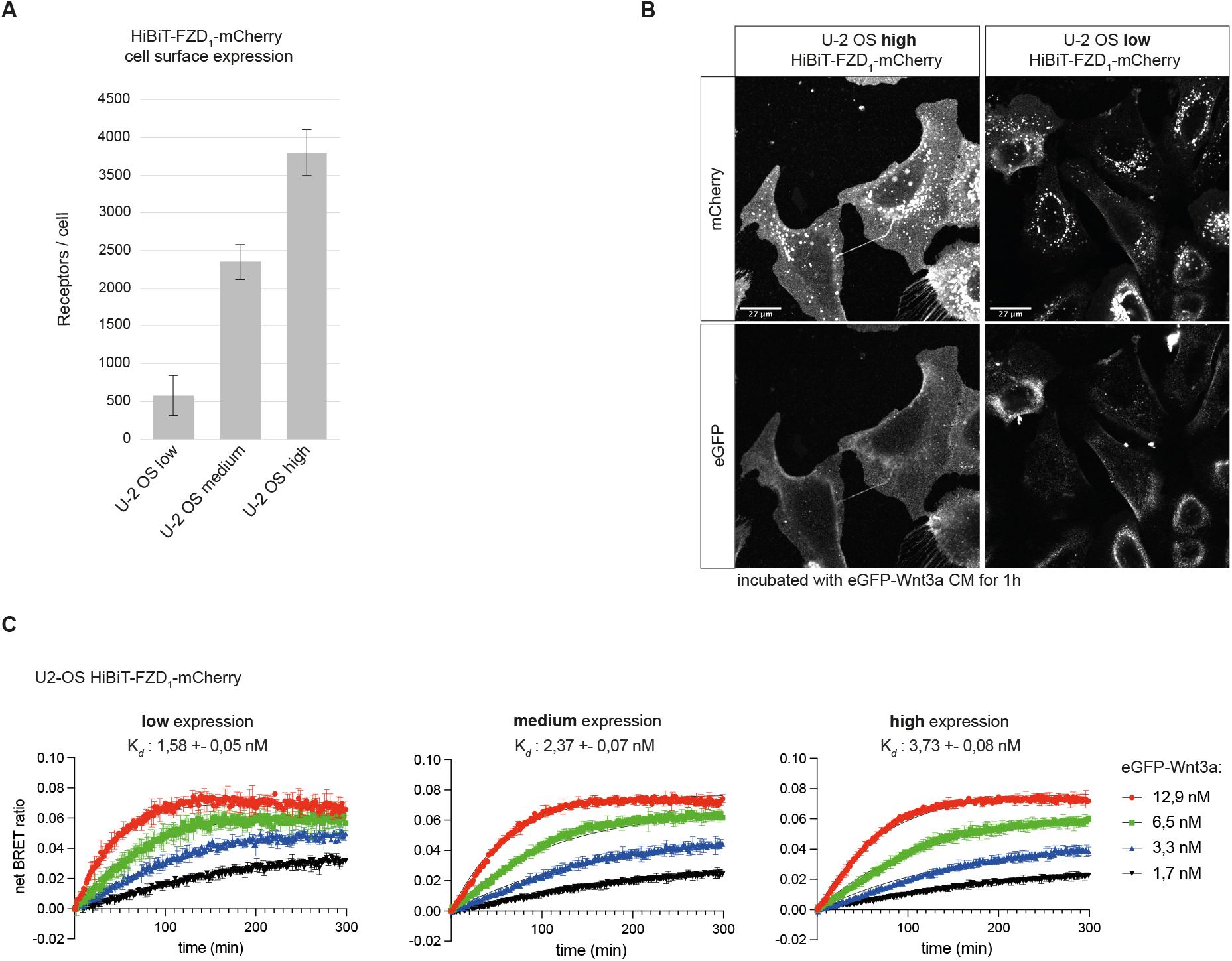
Comparison of U-2 OS stable cell lines with different expression levels of stable integrated of mouse HiBiT-FZD_1_-mCherry gene. **A)** Indirect quantification of HiBiT-FZD_1_-mCherry at the cell surface of FACS sorted U-2 OS stable cell lines expressing three different levels of HiBiT-FZD_1_-mCherry. The average number of cell surface located receptor molecules was estimated using commercially available HiBiT control protein as a reference. **B)** Lazer scanning confocal microscopy image of FACS sorted U-2 OS cells with stable integration of a HiBiT-FZD_1_-mCherry gene, expressed at high and low levels. **C)** Association kinetics of eGFP-Wnt3a binding to HiBiT-FZD_1_-mCherry were determined by NanoBiT/BRET in FACS sorted U-2 OS stable cell lines over a 5 h time period using 1.68, 3.36, 6.48 and 12.96 nM of eGFP-Wnt3a, measured every 78 s. Raw data were fitted to the ‘two or more hot concentrations model’ and are presented as mean ± S.D form n= 3 individual experiments. Three cell lines with different membrane receptor number/cell were used labeled as low, medium high.

### NanoBiT/BRET analysis of Wnt3a, Wnt5a and Wnt16 to HiBiT-FZD_1-10_

For real-time NanoBiT/BRET analysis, which is schematically illustrated in Figure 3A, we FACS sorted a wide range of U-2 OS cell lines expressing low levels of stably integrated HiBiT-FZD_1-10_, as described above (Supplementary Figure 2C, Supplementary Figure 3). The relative levels of HiBiT-FZD’s at the cell surface were estimated from the NanoBiT luminescence generated upon addition of LgBiT and substrate to the cells. Generally, cell surface receptor levels did not vary by more than 2-fold between experiments, however FZD8 and FZD_9_ showed lower, and FZD_10_ higher, levels (Figure 3B). Four concentrations of EGFP-Wnt3a, -5a and -16 were used to generate the association curves needed for calculating kinetic binding affinities. Note that, for some Wnt-FZD combinations, accurate association curves could not be fitted when using either the highest or lowest concentration and one or the other had to be omitted to allow precise calculation of binding affinity (Figure 3C). Of the 30 possible Wnt-FZD combinations for Wnt3a, Wnt5a and Wnt16, binding affinities for 19 pairs could be measured using stable U-2 OS cell lines (Figure 3C). No concentration-dependent increase in BRET ratio was detected for U-2 OS cells expressing either HiBiT-FZD_3_ or HiBiT-FZD_6_, in line with their inability to transduce Wnt/β-catenin signaling (Figure 3, Supplementary Fig. 2D). U-2 OS cells expressing HiBiT-FZD_9_ also failed to show detectable association curves, despite the robust transduction of Wnt/β-catenin signaling (Figure 3, Supplementary Fig. 2D). Indeed, this would fit with the lack of eGFP-Wnt3a binding seen for HiBiT-FZD_9_ expressing U-2 OS cells (Supplementary Fig. 2C).

**Figure 3.**
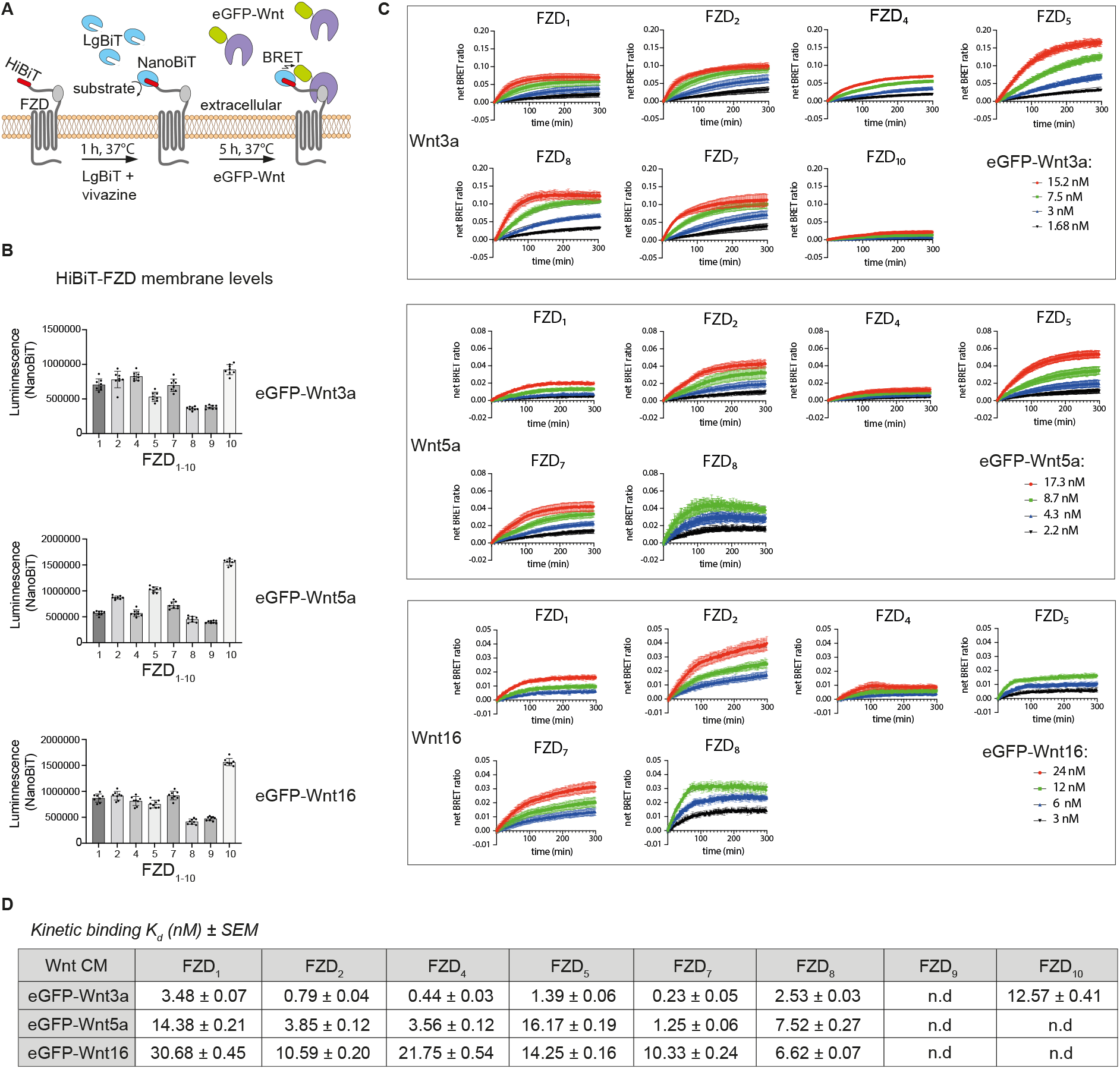
eGFP-Wnt3a, -5a and -16 binding kinetics. **A)** Schematic depiction of NanoBiT/BRET assay. **B)** Estimation of relative HiBiT-FZD cell surface levels in FACS-sorted U-2 OS cells used for each assay. **C)** Association kinetics of eGFP-Wnt3a, eGFP-Wnt5a and eGFP-Wnt16 with U-2 OS cells expressing the indicated HiBiT-FZD proteins. Binding curves were generated over time using 1.5, 3, 7.5,15.2 nM of eGFP-Wnt3a, 2.2, 4.3, 8.7, 17.3 nM of eGFP-Wnt5a or 3, 6, 12, 24 nM of eGFP-Wnt16. BRET ratios were measured every 78 s for a total of 5 h. Raw data were fitted to the ‘two or more hot concentrations model’ and are presented as mean ± SEM form n= 3-4 individual experiments. **D)** Summary of the calculated binding affinities of eGFP-Wnt3a, eGFP-Wnt5a and eGFP-Wnt16 to the various HiBiT-FZD’s. *K*_d_ values are based on data from n=3-4 individual experiments and shown as best-fit value ± SEM. n.d = not detected.

For Wnt3a association to the HiBiT-FZD cell lines, the binding affinity, from highest to lowest was: FZD_7_ > FZD^4^ > FZD_2_ > FZD_5_ > FZD_8_ > FZD_1_ > FZD_10_. This is broadly in agreement with our previous work using HEK293 cells transiently transfected with the same HiBiT-FZD constructs, which express at far higher levels at the cell membrane [21]. Like this study, BRET ratio binding curves for FZD_3_, FZD_6_ and FZD_9_ were mostly undetectable or very low in this previous work [21]. Generally, the k_d_ values calculated here using HiBiT-FZD’s expressed at low level in U-2 OS cells are around 10-fold higher compared to the affinities we calculated previously using transiently transfected HEK293 cells. Although the order of increasing binding affinities was broadly similar between the two studies, one notable exception was FZD_10_, which displays significantly lower affinity here (12.5nM for stable U-2 OS cells compared to 4.3 nM in transiently transfected HEK).

For Wnt5a the FZD binding affinities were, on average, 7-fold lower compared to Wnt3a, although this reduction was greater/less for FZD_5_/FZD_8_, respectively (Figure 3C,D). Similar to eGFP-Wnt3a binding, the strongest association with eGFP-Wnt5a was seen for FZD_7_, followed by FZD_4_ and FZD_2_ (Figure 3D). Indeed, the overall order of binding affinities to FZD’s for Wnt3a and Wnt5a were similar (Wnt3a: FZD_7_ > FZD_4_ > FZD_2_ > FZD_5_ > FZD_8_ > FZD_1_; Wnt5a: FZD_7_ > FZD_4_ > FZD_2_ > FZD_8_ > FZD_1_ > FZD_5_).

Wnt16 had the weakest binding affinities (higher k_d_’s) overall to the HiBiT-FZD cell lines, with FZD_8_, rather than FZD_7_, showing strongest (Figure 3C,D). In contrast to Wnt3a, neither Wnt16 nor Wnt5a generated strong enough binding curves to allow calculation of binding affinities to HiBiT-FZD_10_.

### LRP6 influences the binding affinity between Wnt3a and HiBiT-FZD_1_

Our results demonstrate that NanoBiT/BRET assays can reliably quantify the binding affinities of a wide variety of Wnt/FZD interactions at the cell surface. Wnt proteins however additionally interact with the co-receptor LRP6, forming a trimeric Wnt-FZD-LRP6 complex to transduce Wnt/β-catenin signaling. We therefore used NanoBiT/BRET to study the influence of LRP6 on Wnt/FZD interactions (Figure 4A). To this end we transfected stable U-2 OS HiBit-FZD_1_ cells with either LRP6 or a control pDNA (LacZ) and measured the binding of eGFP-Wnt3a. Compared to FACS sorted U-2 OS cells stably expressing HiBiT-FZD_1_, their transiently transfected control counterparts displayed slightly lower Wnt3a-FZD_1_ binding, with affinities of 3.5 nM and 9.6 nM, respectively (Figure 4B, upper left graph and Figure 3D). Somewhat surprisingly, overexpression of LRP6 resulted in a 3-fold decrease in Wnt3a-FZD_1_ binding affinity (9.6 nM to 27.9 nM) (Figure 4B, compare upper two graphs). We also tested the LRP6 modifier, B3GnT2, which we recently showed can activate Wnt/β-catenin signaling by modifying multiple N-glycans on its extracellular domain [27]. Of note, co-transfection of *LRP6* with *B3GnT2* resulted in a robust (5-fold) increase in Wnt-FZD binding affinity, from 27.9nM to 6.1nM (Figure 4 B, compare graphs on right). A similar, albeit less robust, increase in Wnt-FZD binding affinity was seen upon transfection of *B3GnT2* alone (from 9.6nM to 3.4nM) (Figure 4B, compare graphs on left), suggesting that B3GnT2 may act on endogenous LRP6 and/or FZD to regulate Wnt-FZD association. The amount of HiBiT-FZD_1_ expressed at the cell surface of the U-2 OS cells used in these experiments was similar for all conditions (Figure 4C). Taken together, these results suggest that NanoBiT/BRET analysis of Wnt-FZD interactions is a valuable method for studying the signaling capability of the trimeric Wnt-FZD-LRP6 complex.

**Figure 4.**
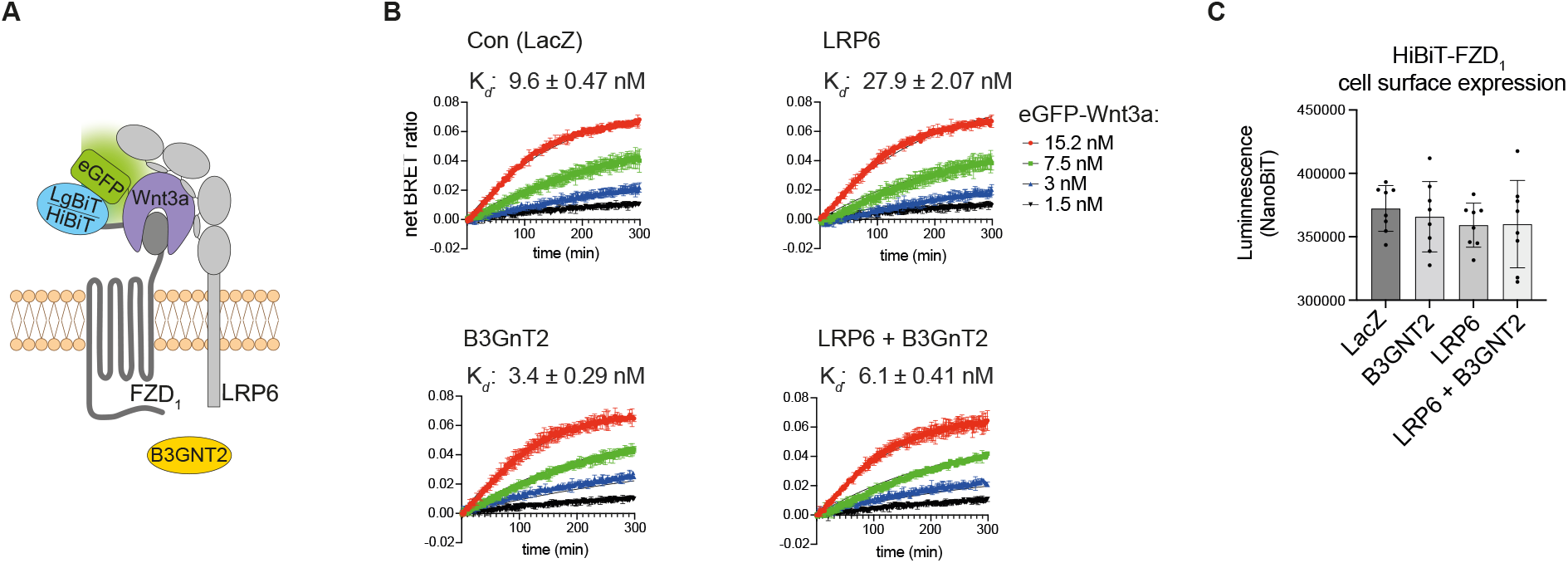
Influence of LRP6 on receptor ligand interaction. **A)** Schematic drawing of trimeric HiBiT-FZD - eGFP-Wnt3a - LRP6 complex formation. **B)** Association kinetics of eGFP-Wnt3a binding to U-2 OS cells stably expressing HiBiT-FZD_1_-mCherry. Cells were transfected with *LacZ, LRP6, B3GNT2* or *LRP6* + *B3GNT2* as indicated and 48 h post transfection the NanoBiT/BRET assay was performed. Association kinetics were determined over time using 1.5, 3, 7.5,15.2 nM of eGFP-Wnt3a CM and measured every 78 s for 5 h. Raw data were fitted to the ‘two or more hot concentrations model’ and are presented as mean ± S.D form n= 3 individual experiments. **C)** Cell surface expression of HiBiT-FZD_1_-mCherry after transfection with the indicated expression plasmids as measured by NanoBiT luminescence.

## Discussion

We continue to develop NanoBiT/BRET as a standard method for real-time quantification of ligand-receptor interactions within the Wnt pathway using full-length proteins in living cells. We have now generated a wide range of cell lines that stably express low levels of HiBiT-FZD_1-10_ and demonstrate that they provide a more accurate system for analysis of Wnt-FZD binding. We have used these cell lines for NanoBiT/BRET analysis together with three different eGFP-Wnt proteins, and we have generated a more comprehensive data set of comparative Wnt-FZD binding affinities. The current study is limited to Wnt3a, Wnt5a and Wnt16 because other fluorescently tagged Wnt proteins could not be clearly shown to bind FZD, or no clear biological activity could be detected for the Wnt CM. Nevertheless, ongoing efforts to expand the availability and use of different fluorescently tagged Wnt’s continues. One encouraging aspect is the apparent ease with which eGFP-tagged Wnt proteins are secreted from cells and obtainable as soluble proteins within the culture medium. The Wnt binding protein Afamin [26] clearly helps the accumulation of eGFP-Wnt proteins in the extracellular medium and thus simplifies the preparation of Wnt conditioned medium, nevertheless its effectiveness varies between Wnt’s. Wnt11 appears to be entirely dependent on Afamin for secretion, others, such as Wnt6, are barely detected in the medium in its absence whereas still others, such as Wnt5a, Wnt8a and Wnt10a appear less dependent. Different mechanisms are reported to account for delivery of Wnt to the extracellular environment [28-31] and specific Wnt’s may have mechanistic preferences, perhaps explaining the differences in how Afamin affected our results.

It is informative to compare the work presented here using the stable U-2 OS cell lines with our previous study using transient transfection of HEK293 cells, which produces significantly higher receptor levels [21]. On average, we see a near 10-fold increase in Wnt-FZD binding affinities using the stable U-2 OS cell lines expressing low HiBiT-FZD levels. This fits well with our hypothesis that accurate and precise quantification of ligand-receptor interactions is best achieved when signalling proteins interact with receptors at the cell surface under physiologically relevant conditions. Fortunately, there appears to be a range of such physiological relevant “low” expression levels because stable cell lines expressing significantly higher receptor levels compared to endogenously tagged receptors exhibit similar binding affinities [23]. Only transiently transfected cells that express far higher receptor levels have significantly lower binding affinities. Thus, we can generate and confidently use stable cell lines with relatively low levels of tagged receptors and do not require CRISPR-mediated endogenous tagging.

In contrast to the strong differences in Wnt3a-FZD binding affinities between transiently transfected HEK293 cells and U-2 OS stable cell lines, the relative order of binding affinities between the two studies remained similar, apart from FZD_10_. This FZD however also presented relatively low BRET ratio values in spite of showing the highest cell surface expression levels (Figure 3B). In summary, the relative consistency between our studies using different cells indicates that NanoBiT/BRET binding should be suitable for a variety of cells, however future studies using a greater variety of cell types is needed to confirm this.

Despite the markedly lower overall binding strengths to FZD’s for Wnt5a compared to Wnt3a, one somewhat surprising finding was the similarity for Wnt3a and Wnt5a in their relative order of binding affinities to the different FZD’s. This would point to the nature of binding between Wnt/FZD being mostly attributable to the more conserved elements of Wnt’s. Nevertheless, Wnt16 does show some striking differences, such as FZD_8_ being the strongest binding partner and FZD_4_ one of the weakest, which is in contrast to the binding profiles of either Wnt3a or Wnt5a.

In our experiments using HiBiT-FZD_1_ expressing cells with/without LRP6 co-expression, LRP6 appears to regulate the binding strengths between Wnt and FZD within the Wnt-FZD-LRP6 complex. This may provide some additional insight with respect to the mechanisms governing Wnt signalosome formation. Evidence exists that one Wnt molecule can associate with more than one FZD-CRD unit [32]. It is tempting to hypothesise that LRP6 can influence the ability of Wnt to form different stoichiometries with FZD’s and future work using tools such as NanoBiT/BRET-based ligand-receptor interactions should help provide answers. Additionally, our results demonstrate that enhanced glycosylation of the extracellular region of LRP6 by B3GnT2 also influences Wnt-FZD association. It is noteworthy that B3GnT2 extends polylactosamine chains close to potential Wnt binding sites on LRP6 and may therefore sterically alter Wnt-FZD interactions. This may indeed partially account for B3GnT2’s ability to enhance Wnt/LRP6 signalling [27].

Our strategic use of NanoBiT/BRET methodology to accurately quantify the large repertoire of ligand-receptor interactions within the Wnt signalling pathway appears to be a valid approach. The generation of a wide range of cell lines expressing physiologically relevant levels of HiBiT-FZD has allowed us, for the first time, to assemble a more global comparative overview as well as accurate quantification of Wnt-FZD interactions on cell membranes.

## Materials and Methods

### Cell Culture and Ligands

Human embryonic kidney 293T (HEK293T) cells (CLS catalog number 300189), ΔFZD_1-10_ HEK293 cells [22], U-2 OS cells (ATCC^®^ HTB-96™) and U-2 OS cell lines with stable integration of mouse HiBiT-FZD_1,2,4,5,7,8,9,10_-mCherry (generated in this project) were cultured in Dulbecco’s modified Eagle medium (Gibco, Thermo Fisher, Waltham, MA) supplemented with 10% fetal bovine serum (FBS, Thermo Fisher) and 1% penicillin-streptomycin (P/S; Gibco; Thermo Fisher). NCI-H1703 with stable integration of mouse HiBiT-FZD_1,4,5,8,10_-mCherry (generated in this project) were cultured in Roswell Park Memorial Institute (RPMI) 1640 medium (Gibco, Thermo Fisher, Waltham, MA) supplemented with 10% fetal bovine serum, 1 mM sodium pyruvate (Gibco, Thermo Fisher), 1% P/S. HTC116 cells with stable integration of mouse HiBiT-FZD1-mCherry (generated in this project) were cultured in McCoy’s 5a modivied medium (Gibco, Thermo Fisher) supplemented with 10% fetal bovine serum (FBS, Thermo Fisher) and 1% penicillin-streptomycin (P/S; Gibco; Thermo Fisher). All cell lines were maintained at 37°C and 5% CO_2_. Expi293™ suspension cells (Thermo Fisher, A14527) were cultured in Expi293™ expression medium (Thermo Fisher) at 37°C and 8% CO2 with 125 rpm orbital shaking in a New Brunswick S41iCO2 shaking incubator (Eppendorf). Cell densities and viability were determined using a Countess II automated cell counter (Life Technologies).

### Preparation of eGFP-Wnt3a, eGFP-Wnt5a and eGFP-Wnt16 CM in Expi293™ suspension cells

Expi293™ suspension cells growing in Expi293™ expression medium (60 ml, 2.5 × 10^6^ cells/ml) were co-transfected with 10 µg of either pCS2+-eGFP-Wnt3a, pCS2+-eGFP-Wnt5a or pCS2+-eGFP-Wnt16 each together with 50 µg of pCMV-His-Afamin plasmid using ScreenFect® UP-293 (ScreenFect GmbH) according to the manufacturer’s instructions. eGFP-Wnt CM were collected 96 h post transfection. The corresponding control CM was generated from cells transfected with pCS2+ plasmid. For downstream experiments, the “raw” CM was 5-fold concentrated using Vivaspin turbo 30,000-molecular-weight-cutoff ultra filters (Satorius AG) and exchanged to the desired cell culture medium using Sephadex G-25 PD10 desalting columns (GE Healthcare Bio-Science). The final concentration and integrity of eGFP-Wnt-3a in the CM samples were determined using ELISA (GFP ELISA® kit, Abcam, ab171581).

### Generation of eGFP-Wnt CM in HEK293T cells

1,2 ^*^10^6^ HEK293T cells in 6 well were co-transfected with 0,7 µg pCMV3-His-hAfamin, 1 µg pCS2+ together with 0.3 µg pCS2+ eGFP-Wnt fusion construct for Wnt1, Wnt2b, Wnt3a, Wnt5a, Wnt6, Wnt7a, Wnt8a, Wnt9a, Wnt10a, Wnt10b, Wnt11 and Wnt16, or co-transfected with 1,7 µg pCS2+ and 0,3 µg pCS2+ eGFP-Wnt fusion construct using ScreenFect^®^ A (ScreenFect GmbH) according to the manufacturer’s 1 step protocol. 72 h post transfection medium was discarded, 2 ml fresh medium was added to the cells and incubated for further 72 h before harvest. Conditioned medium (CM) was harvested and centrifuged for 5 min at 1500 x g to pellet possible dead cells. 1.5 ml CM was mixed with 75 µl of 20% Triton-X 100 to a final concentration of 1% Triton-X 100. Cibacron blue 3G coupled to Sepharose 6 Fast Flow (Blue-Sepharose 6 Fast Flow, GE Healthcare) beads were washed three times with Blue-Sepharose buffer (BS-buffer) (150 mM KCl, 50 mM Tris-HCl, pH 7.5, 1% Triton X-100) containing Complete® protease inhibitor mixture (Roche). Washed beads were added to the eGFP-Wnt CM and rotated overnight at 4 °C. The next day, beads were washed three times in BS-buffer, Laemmli sample buffer was added, and samples were heat denaturated. SDS-PAGE/ Western blotting was performed and secreted eGFP-Wnt proteins were detected using anti GFP antibody (Santa Cruze Biotechnology, sc-9996).

### Plasmids

NFAT as well as AP-1 Luciferase reporter were generated by replacing TCF/LEF binding sites of M50 Super 8xTOPFlash (a gift from Randall Moon, Addgene plasmid 12456) by NFAT or AP-1 response elements. For NFAT, the binding site from human interleukin 2 (IL-2) gene (Mattila et al., 1990) was ordered as complementary DNA oligos flanked by KpnI and XhoI sites (NFAT binding site, 5’ -AAC TCG AGC GCC TTC TGT ATG AAA CAG TTT TTC CTC CGG TAC CAA A -3’). The oligos were annealed and inserted between KpnI and XhoI restriction sites of M50 plasmid. For AP-1 Luciferase reporter, the AP-1 response element of three repeats of two alternative AP-1 binding sites [33] were ordered as oligos flanked by KpnI and NheI restriction sites (AP-1 binding site, 5’ -AAG GTA CCT GAG TCA GTG ACT CAG TGA GTC AGT GAC TCA GTG AGT CAG TGA CTC AGC TCG AGA AA -3’). The oligos were annealed and inserted between KpnI and NheI restriction sites of M50 plasmid. pCMV3-His-hAfamin was obtained from Sino Biological (HG13231-CH), pCS2+ eGFP-Wnt-3a (Wesslowski et al., 2020). N-terminal eGFP-(GGSG)-Wnt fusion proteins for Wnt1, Wnt2b, Wnt5a, Wnt6, Wnt7a, Wnt8a, Wnt9a, Wnt10a, Wnt10b, Wnt11 and Wnt16 were generated by inserting the synthetic human codon optimized ORF of eGFP-Wnt DNA fragment (Invitrogen GeneArt Strings DNA Frgaments, Thermo Fisher Scientific) into pCS2+ expression vector using BamHI and XbaI restriction sites. To generate pEF1*α* HiBiT-FZD_1,2,4,6,7,8,9,10_-mCherry-IRES-NEO, FZD_1,2,4,6,7,8,9,10_-mCherry ORFs without signal peptides were PCR amplified from the corresponding pmCherry-FZD-mCherry plasmids [22] flanked by XbaI and NotI restriction sites. 5-HT_3_A signal peptide followed upstream by HiBiT sequence [22] and flanked by XhoI and XbaI restriction sites was ordered as complementary DNA oligos (5’ TCG AGG CCA CCA TGC GGC TCT GCA TCC CGC AGG TGC TGT TGG CCT TGT TCC TTT CCA TGC TGA CAG GGC CGG GAG AAG GCA GCC GGG TGA GCG GCT GGC GGC TGT TCA AGA AGA TTA GCT -3’ (forward) and 5’ CTA GAG CTA ATC TTC TTG AAC AGC CGC CAG CCG CTC ACC CGG CTG CCT TCT CCC GGC CCT GTC AGC ATG GAA AGG AAC AAG GCC AAC AGC ACC TGC GGG ATG CAG AGC CGC ATG GTG GCC-3’ (reverse)). The oligos were annealed and together with the amplified FZD1,2,4,6,7,8,9,10-mCherry ORFs inserted between XhoI and NotI restriction site of pEF1*α*-IRES-NEO (a gift from Thomas Zwaka, Addgene plasmid 28019).

### Generation of cell lines stably expressing HiBiT-FZD-mCherry

NCI-1703, HTC116 or U-2 OS cells were transfected with 2µg pEF1a-HiBiT-FZD-mCherry-IRES-Neo in 6 well plates using ScreenFect^®^ A (ScreenFect GmbH) according to the manufacturer’s 1 step protocol. Twenty-four hours post transfection, the medium was exchanged, and 48 h post transfection cells were transferred to 10-cm^2^ dishes and cultured in their respective cell culture medium supplemented with 1 mg/ml G418 (Sigma-Aldrich). Selection was performed for 7 days. Selected cells were FACS sorted according to cell surface receptor expression.

### Fluorescence-activated cell sorting (FACS)

FACS was performed with a FACSAriaTM Flow Cytometer (BD Biosciences). In order to sort U-2 OS for stable integration of HiBiT-FZD-mCherry gene with cell surface located HiBiT-FZD-mCherry, cells were detached using 5 mM EDTA (Roth) in PBS^-/-^ (Gibco, Thermo Fisher) and collected in DMEM (Gibco) supplemented with 10% FBS (Gibco). A total amount of approximately 1 × 107 cells was used for sorting. Cells were resuspended in 1 ml ice cold FACS buffer (2% FBS; 2 mM EDTA; PBS^-/-^). Surface HiBiT-FZD-Cherry was labeled with 1µg/ml anti-HiBiT antibody (#N7200 Promega) in FACS buffer for 45 min on ice. Cells were washed two times with 5 ml ice cold FACS buffer and labeled with 2,5 µg/ml anti mouse Alexa 488 (A21204, Invitrogen, Thermo Fisher) for 30 min on ice. Cells were washed two times with 5 ml FACS buffer and resuspended in 4 ml FACS buffer for sorting. Gating for low, medium, high cell surface protein signal was set in BD Diva software and cells were single cell sorted in 96 well plates. 5 weeks after sorting clonal lines were re-analyzed with FACS for their cell surface expression levels and verified via FACS analysis.

### Reporter Gene Assay

To test the biological activity of eGFP-Wnt-3a, eGFP-Wnt-5a and eGFP-Wnt-16, 5,5 × 10^4^ HEK293T cells cultured in 96-well plates were transfected with 20 ng TCF firefly luciferase (TOP-FLASH), 2 ng CMV Renilla luciferase, and 78 ng pCS2+ empty plasmid using ScreenFect® A according to the manufacturer’s 1-step protocol (ScreenFect GmbH). Twenty-four hours post transfection, medium was replaced by control, eGFP-Wnt-3a CM, eGFP-Wnt-5a CM or eGFP-Wnt-16 CM and cells were incubated for another 24 h before harvesting cell lysates.

For the luciferase assays, cells from 96-well plates were harvested in 43 µl 1x Passive Lysis Buffer (Promega) and processed according to the manufacturer’s protocol. TOPFLASH luciferase values were normalized to control Renilla luciferase values. All error bars shown are standard deviations from the mean (± S.D.) of the indicated number of *n* = 4 independent biological samples within an experiment, and all experiments were performed at least 3 times, unless indicated otherwise in the figure legends. Data were analyzed using GraphPad Prism 8.

### WB analysis

eGFP-Wnt-3a, eGFP-Wnt-5a and eGFP-Wnt-16 were mixed with Laemmli sample buffer and heat denatured. Samples were separated by SDS-PAGE before transfer to a PVDF membrane using a Bio-Rad Transblot-Turbo system (Bio-Rad). Membranes were blocked at room temperature for 1 h in 5% BSA–TBST blocking buffer (5% BSA, 137 mM NaCl, 2.7 mM KCl, 19 mM Tris base [pH 7.4], 0.1% Tween-20) and transferred to a BioLane HTI automated Western blotting processor for antibody incubation and washing steps. The following antibodies were used: anti anti-GFP (ab1828, 1:2,000, Abcam) and HRP-conjugated anti-rabbit or anti-mouse secondary antibodies (Dako). For semiquantitative detection of protein bands, the membranes were incubated with ECL Prime (GE-Healthcare Bio-science) and imaged using a ChemiDocTM touch imaging system (Bio-Rad).

### NanoBiT/BRET Binding assay

U-2 OS cells stably expressing HiBiT-FZD-mCherry were seeded as 6500 cells/well onto a poly-D-lysine (Gibco, Thermo Fisher) -coated white 96-well cell culture plate with clear flat bottom (VWR part of avantor). Twenty-four hours later, the cells were washed once with 200 µl non-phenol red DMEM (Gibco, Thermo Fisher) supplemented with 5% FBS, 10 mM HEPES (Gibco, Thermo Fisher). The cells were preincubated with 50 µl non-phenol red DMEM supplemented with Vivazine (1:100 dilution; Promega), LgBiT (1:100 dilution, Promega), 5% FBS and 10 mM HEPES for 1 h at 37 °C without CO_2_. Subsequently, 50 µl of four different concentrations of eGFP-Wnt-3a (1.68, 3, 7.5 and 15.2 nM), eGFP-Wnt-5a (2.2, 4.3, 8.7, 17.3 nM), eGFP-Wnt-16 (3, 6, 12, 24 nM) conditioned medium or control conditioned medium supplemented with Vivazine (1:150 dilution, Promega), 5% FBS, 10 mM HEPES was added, and the BRET signal was measured every 80 s for 300 min at 37 °C.

For NanoBiT/BRET assay with stable HiBiT-FZD_1_-mCherry U-2 OS cells co-expressing B3GNT2 and LRP6, a total of 10,000 cells were seeded in poly-d-lysine (Gibco, Thermo Fisher)-coated white 96-well cell culture plates with clear bottom. After 24 hours cells were transfected with lacZ (control up to 100 ng), B3GnT2 (1 ng), LRP6 (20 ng), MESD (5 ng) using Lipofectamine 3000 Transfection Reagent (Thermo Fisher). Next day cells were washed once with 200 μL of non-phenol red DMEM supplemented with 10 mM HEPES and 5% FBS. The cells were preincubated with 50 μL of a mix of Vivazine (1:50 dilution) and LgBiT (1:100 dilution) in a complete, non-phenol red DMEM supplemented with 10 mM HEPES for 1 h at 37 °C without CO_2_. Subsequently, 50ul of four different concentrations (1.5, 3, 7.5 and 15.2 nM) of eGFP-WNT-3A conditioned medium supplemented with 5% FBS and 10 mm HEPES was added, and the BRET signal was measured every 80 s for 300 min at 37 °C.

eGFP fluorescence was measured prior to reading BRET (excitation, 470–15 nm; emission, 515–20 nm). A with sticker (VWR) was sticked under the plate and the BRET ratio was determined as the ratio of light emitted by eGFP-tagged ligands (energy acceptor) and light emitted by HiBiT-tagged receptors (energy donors). The BRET acceptor (bandpass filter, 535– 30 nm) and BRET donor (bandpass filter, 450–80 nm) emission signals were measured using a CLARIOstar microplate reader (BMG). Data were analyzed using GraphPad Prism 8.

### Estimation of receptor density

Stable U-2 OS HiBiT-FZD_1_-mCherry cells were seeded in a white 96 well plate with transparent bottom and 48h later the measurement was started. To prepare a standard curve, 15 samples HiBiT control protein (Promega, #N3010) were prepared ranging from 1 fM to 100 mM in non-phenol red DMEM supplemented with 5% FBS, 10mM HEPES and 90 µl / well each was distributed as duplicates in white 96 well plates with transparent bottom. Cells were washed once with 200 µl non-phenol red DMEM and covered with 90 µl non-phenol red DMEM supplemented with 5% FBS and 10mM HEPES. Reaction mix was prepared by supplementing non-phenol red DMEM with LgBiT (1:20 Promega) and Furimazine (1:10, Promega). 10 µl of the reaction mix was added to each well, carefully mixed and incubated for 10 min at 37°C without CO_2_. A white sticker was sticked to the bottom of the plate and the luminescence emitted by the complemented NanoBiT Luciferase was measured (460-500 nm, 200ms integration time). Ater measurement cells were detached and counted using a Countess II automated cell counter (Life Technologies). Luminescence/cell was calculated and together with the standard curve and the molecular weight of the HiBiT control protein the receptor number/cell was calculated.

### Data Analysis and Statistics

All data were analyzed in GraphPad Prism 8 (San Diego, CA, USA) using built-in equations. All data presented in this study come from *n* individual experiments (at least three biological replicates) with each individual experiment performed typically in duplicates for each tested condition. Data points on the binding curves represent mean ± SEM. Kinetic binding data were analyzed using the association model with two or more hot ligand concentrations. Binding affinity values (*K*_d_) are presented as a best-fit *K*_d_ with SEM.

### Confocal laser scanning microscopy

For the microscopy analysis of eGFP-Wnt fusion proteins binding to Frizzled receptors, NCI-1703 cells with stable integration of HiBiT-FZD_4,5,8,10_-mCherry gene were seeded in µ-Slide 18-well chambers (Ibidi, catalog no. 81816) so that they were 80% confluent for microscopy after 2 days. Prior imaging the individual eGFP-Wnt CM for Wnt1, Wnt2b, Wnt3a, Wnt5a, Wnt6, Wnt7a, Wnz8a, Wnt9a, Wnt10a, Wnt10b, Wnt11, Wnt16 were added to the cells and incubated for 1 h. For microscopy, cell medium was exchanged to a mix of 50% FluoroBrite DMEM (Gibco, Thermo Fisher) and 50% non-phenolred RPMI (Gibco, Thermo Fisher) supplemented with 10% FBS, and 10 mM HEPES (Gibco, Thermo Fischer). For microscopy analysis of eGFP-Wnt3a binding to Frizzled recepors in U-2 OS cells. U-2 OS cells with stable integration of HiBiT-FZD_1,2,3,4,5,6,7,8,9,10_-mCherry were seeded in µ-Slide 18-well chambers (Ibidi, catalog no. 81816) so that they were 80% confluent for microscopy after 2 days. Prior imaging eGFP-Wnt3a CM was added to the cells and incubated for 3 h. For microscopy, cell medium was exchanged to FluoroBrite DMEM (Gibco, Thermo Fisher) supplemented with 10% FBS, and 10 mM HEPES (Gibco, Thermo Fischer). Image acquisition was performed using a Zeiss LSM 800 microscope (Zeiss, Jena, Germany) fitted with a 40x/1.2 oil differential interference contrast (UV) VIS-IR Plan-Apochromat objective (Zeiss, Jena, Germany) and a GaAsP-PMT detector. The red mCherry and green eGFP fluorescent proteins were excited at 561 nm and 488 nm, respectively, and their respective emissions were captured in the range of 570–700 nm and 400–576 nm, employing the standard filter sets where appropriate. Images were analyzed using Fiji [34].

## Acknowledgements

We thank Christine Blattner (KIT, Karlsruhe) for providing the U-2 OS cells. This work was funded by the Deutsche Forschungsgemeinschaft (DFG, German Research Foundation) – project number 331351713 – SFB 1324 (project A06 to G.Davidson).

## Author Contributions

G.D. and J.W. conceived and supervised the project, its planning and execution. J.W. performed all experiments apart from FACS sorting. S.S. provided help with LRP6/B3GnT2 experiments. Me.R. performed all FACS experiments. Mi.R. prepared Wnt CM. and provided general help with experiments. All co-authors contributed to the preparation of Figures. G.D. wrote the manuscript with input from J.W.

## Competing interests

The authors declare no conflict of interest.

## Notes

### Competing Interest Statement

The authors have declared no competing interest.

## References

1. Niehrs, C., The complex world of WNT receptor signalling. Nat Rev Mol Cell Biol, 2012. 13(12): p. 767–79.

2. Wiese, K.E., R. Nusse, and R. van Amerongen, Wnt signalling: conquering complexity. Development, 2018. 145(12).

3. Holstein, T.W., The evolution of the Wnt pathway. Cold Spring Harb Perspect Biol, 2012. 4(7): p. a007922.

4. Holzem, M., M. Boutros, and T.W. Holstein, The origin and evolution of Wnt signalling. Nat Rev Genet, 2024. 25(7): p. 500–512.

5. Endo, M., K. Kamizaki, and Y. Minami, The Ror-Family Receptors in Development, Tissue Regeneration and Age-Related Disease. Front Cell Dev Biol, 2022. 10: p. 891763.

6. Davidson, G., LRPs in WNT Signalling. Handb Exp Pharmacol, 2021. 269: p. 45–73.

7. Rim, E.Y., H. Clevers, and R. Nusse, The Wnt Pathway: From Signaling Mechanisms to Synthetic Modulators. Annu Rev Biochem, 2022. 91: p. 571–598.

8. Rulifson, E.J., C.H. Wu, and R. Nusse, Pathway specificity by the bifunctional receptor frizzled is determined by affinity for wingless. Mol Cell, 2000. 6(1): p. 117–26.

9. Bhanot, P., et al., A new member of the frizzled family from Drosophila functions as a Wingless receptor. Nature, 1996. 382(6588): p. 225–30.

10. Kikuchi, A., et al., New insights into the mechanism of Wnt signaling pathway activation. Int Rev Cell Mol Biol, 2011. 291: p. 21–71.

11. Kikuchi, A., et al., Wnt5a: its signalling, functions and implication in diseases. Acta Physiol (Oxf), 2012. 204(1): p. 17–33.

12. Mikels, A.J. and R. Nusse, Purified Wnt5a protein activates or inhibits beta-catenin-TCF signaling depending on receptor context. PLoS Biol, 2006. 4(4): p. e115.

13. Acebron, S.P. and C. Niehrs, beta-Catenin-Independent Roles of Wnt/LRP6 Signaling. Trends Cell Biol, 2016.

14. Koca, Y., G.M. Collu, and M. Mlodzik, Wnt-frizzled planar cell polarity signaling in the regulation of cell motility. Curr Top Dev Biol, 2022. 150: p. 255–297.

15. Habib, S.J. and S.P. Acebron, Wnt signalling in cell division: from mechanisms to tissue engineering. Trends Cell Biol, 2022. 32(12): p. 1035–1048.

16. Tejeda-Munoz, N. and E.M. De Robertis, Wnt, GSK3, and Macropinocytosis. Subcell Biochem, 2022. 98: p. 169–187.

17. Niehrs, C., Function and biological roles of the Dickkopf family of Wnt modulators. Oncogene, 2006. 25(57): p. 7469–81.

18. He, X., et al., LDL receptor-related proteins 5 and 6 in Wnt/beta-catenin signaling: arrows point the way. Development, 2004. 131(8): p. 1663–77.

19. Acebron, S.P., et al., Mitotic wnt signaling promotes protein stabilization and regulates cell size. Mol Cell, 2014. 54(4): p. 663–74.

20. Gratz, L., et al., NanoBiT- and NanoBiT/BRET-based assays allow the analysis of binding kinetics of Wnt-3a to endogenous Frizzled 7 in a colorectal cancer model. Br J Pharmacol, 2024. 181(20): p. 3819–3835.

21. Kozielewicz, P., et al., Quantitative Profiling of WNT-3A Binding to All Human Frizzled Paralogues in HEK293 Cells by NanoBiT/BRET Assessments. ACS Pharmacol Transl Sci, 2021. 4(3): p. 1235–1245.

22. Wesslowski, J., et al., eGFP-tagged Wnt-3a enables functional analysis of Wnt trafficking and signaling and kinetic assessment of Wnt binding to full-length Frizzled. J Biol Chem, 2020. 295(26): p. 8759–8774.

23. Eckert, A.F., et al., Measuring ligand-cell surface receptor affinities with axial line-scanning fluorescence correlation spectroscopy. Elife, 2020. 9.

24. Willert, K., et al., Wnt proteins are lipid-modified and can act as stem cell growth factors. Nature, 2003. 423(6938): p. 448–52.

25. Takada, R., et al., Assembly of protein complexes restricts diffusion of Wnt3a proteins. Commun Biol, 2018. 1: p. 165.

26. Mihara, E., et al., Active and water-soluble form of lipidated Wnt protein is maintained by a serum glycoprotein afamin/alpha-albumin. Elife, 2016. 5.

27. Xu, R., et al., N-Glycosylation of LRP6 by B3GnT2 Promotes Wnt/beta-Catenin Signalling. Cells, 2023. 12(6).

28. Mittermeier, L. and D.M. Virshup, An itch for things remote: The journey of Wnts. Curr Top Dev Biol, 2022. 150: p. 91–128.

29. Takada, S., et al., Differences in the secretion and transport of Wnt proteins. J Biochem, 2017. 161(1): p. 1–7.

30. Zhang, L. and J.L. Wrana, The emerging role of exosomes in Wnt secretion and transport. Curr Opin Genet Dev, 2014. 27: p. 14–9.

31. Bartscherer, K. and M. Boutros, Regulation of Wnt protein secretion and its role in gradient formation. EMBO Rep, 2008. 9(10): p. 977–82.

32. DeBruine, Z.J., H.E. Xu, and K. Melcher, Assembly and architecture of the Wnt/beta-catenin signalosome at the membrane. Br J Pharmacol, 2017. 174(24): p. 4564–4574.

33. Angel, P., et al., Phorbol ester-inducible genes contain a common cis element recognized by a TPA-modulated trans-acting factor. Cell, 1987. 49(6): p. 729–39.

34. Schindelin, J., et al., Fiji: an open-source platform for biological-image analysis. Nat Methods, 2012. 9(7): p. 676–82.

