## Supplementary Figures for "Quantification of Wnt3a, Wnt5a and Wnt16 Binding to Multiple Frizzleds Under Physiological Conditions using NanoBit/BRET"

Supplementary Figures for Wesslowski et al. 2025

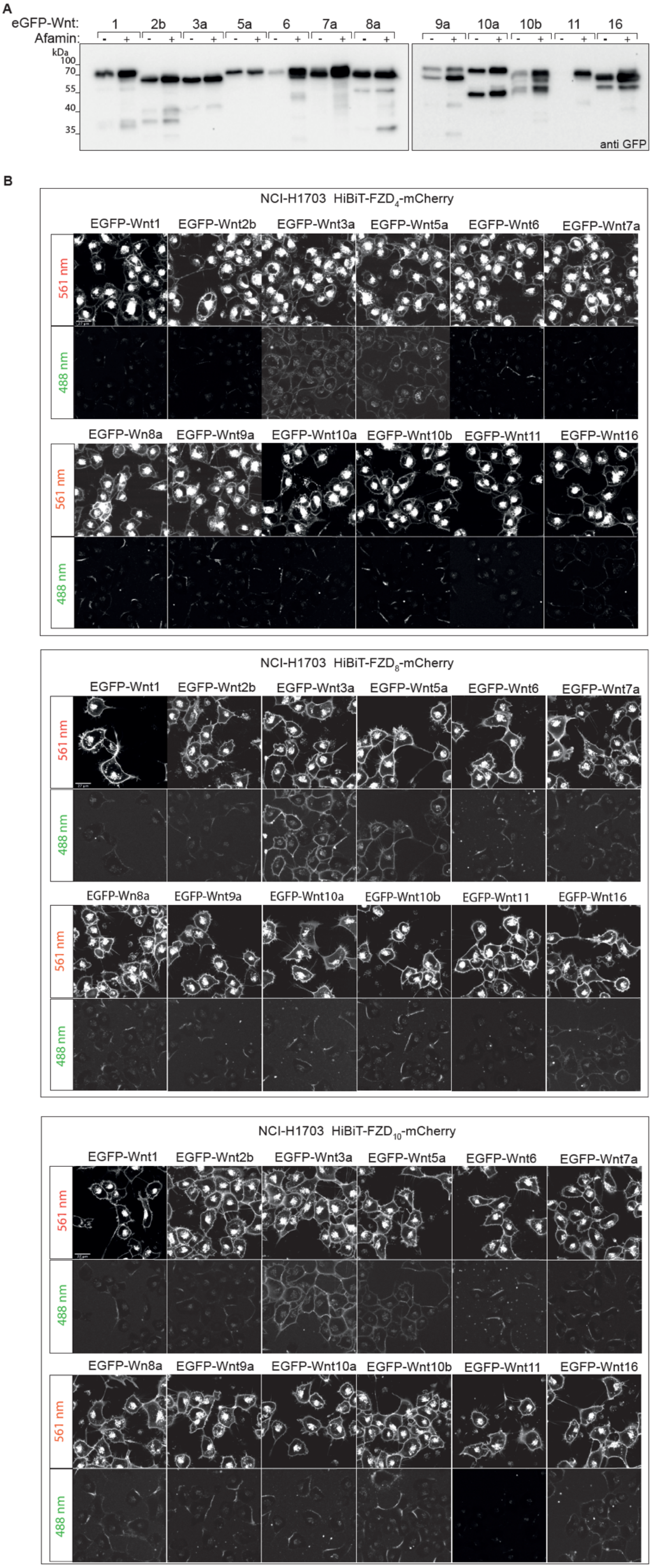


***Supplementary Figure 1. Generation and testing of eGFP-Wnt fusion proteins. A)*** *Western Blot (WB) of conditioned medium (CM) harvested from HEK293T cells transfected with the indicated constructs, confirming the secretion of full length eGFP-Wnt proteins with/without Afamin co-expression.* ***B)*** *Laser scanning confocal microscopy images of NCI-H1703 cells with stable integration of mouse HiBiT-FZD$_{4,8,10}$-mCherry, incubated for 3 h with the indicated eGFP-Wnt CM derived from Expi293$^{TM}$ suspension cells.*

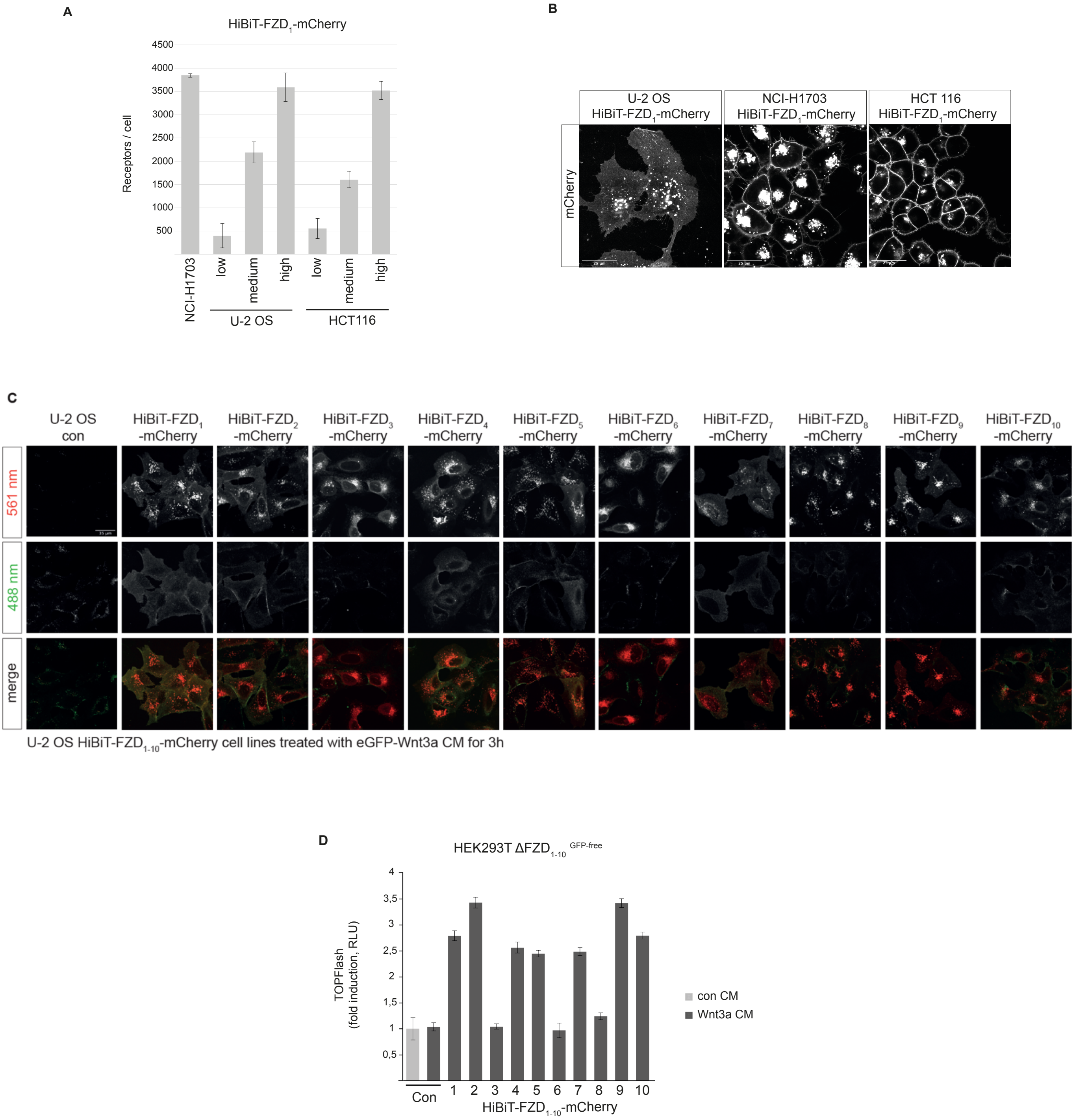


**Supplementary Figure 2. Comparison of stable expression levels of HiBiT-FZD$_1$ in NCI-1703 U-2 OS and HCT116 cells. A)** Indirect quantification of HiBiT-FZD$_1$-mCherry at the cell surface of the indicated FACS sorted stable cell lines. **B)** Lazer scanning confocal microscopy images comparing the expression of HiBiT-FZD$_1$-mCherry in the indicated FACS sorted cells. **C)** Laser scanning confocal microscopy images of living U-2 OS cells with stable integration of mouse HiBiT-FZD-mCherry constructs as indicated. Cells were incubated for 3 h with eGFP-Wnt3a condition medium (CM) derived from Expi293$^{TM}$ suspension cells. **B)** TOPFlash reporter assay in ΔFZD$_{1\text{-}10}^{GFP\text{-}free}$ HEK293 cells, showing the relative Wnt/β-catenin signaling activity of the indicated mouse HiBiT-FZD-mCherry receptors. Error bars represent ± S.D. form means of 4 independent biological samples.

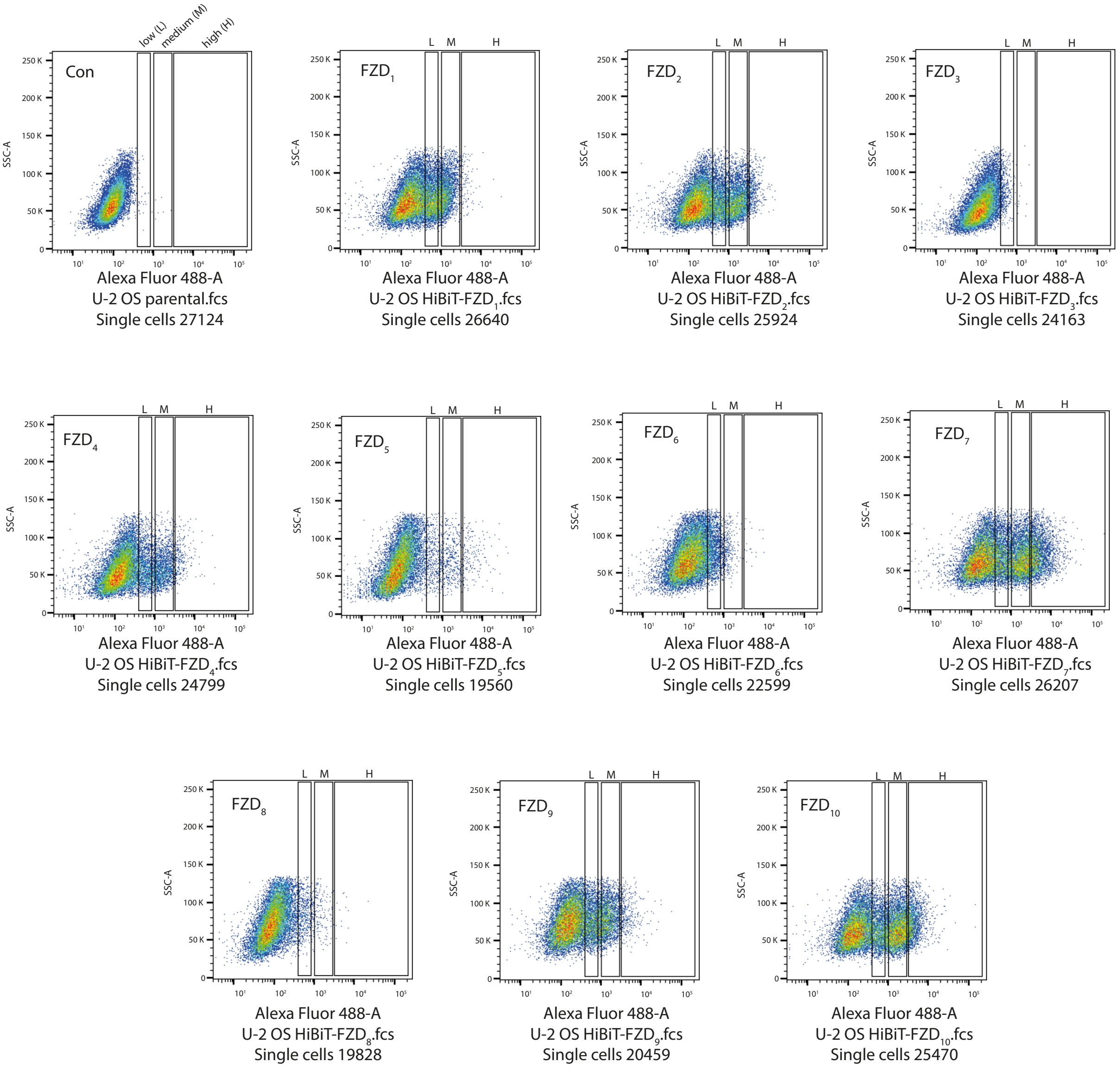


**Supplementary Figure 3. FACS sorting of U-2 OS cells stably expressing HiBiT-FZD$_{1\text{-}10}$.** Single cell sorting of U-2 OS cells stably expressing HiBiT-FZD-mCherry at the cell surface. U-2 OS cell suspensions were incubated with mouse anti-HiBiT antibody following by labelling with anti-mouse Alexa 488 antibody. All cells were analyzed on one dimensional plots of the green channel- 488 nm for HiBiT positive cells. Gated in black as low (L), medium (M) and high (H) are cell populations with di erent expression levels of HiBiT-FZD at the cell surface. Parental U-2 OS cells were used as background control. Cells gated for low, medium and high were single cell sorted in 96 well plate.
